# Adding function to the genome of African *Salmonella* ST313

**DOI:** 10.1101/450098

**Authors:** Rocío Canals, Disa L. Hammarlöf, Carsten Kröger, Siân V. Owen, Wai Yee Fong, Lizeth Lacharme-Lora, Xiaojun Zhu, Nicolas Wenner, Sarah E. Carden, Jared Honeycutt, Denise M. Monack, Robert A. Kingsley, Philip Brownridge, Roy R. Chaudhuri, Will P. M. Rowe, Alexander V. Predeus, Karsten Hokamp, Melita A. Gordon, Jay C. D. Hinton

**Author notes:** Disa L. Hammarlöf, Department of Cell and Molecular Biology, Uppsala University, Uppsala, 751 24, Sweden. Carsten Kröger, Department of Microbiology, School of Genetics and Microbiology, Moyne Institute of Preventive Medicine, Trinity College Dublin, Dublin 2, Ireland. Siân V. Owen, Department of Biomedical Informatics, Harvard Medical School, Boston, Massachusetts, 02115, USA. Will P. M. Rowe, Scientific Computing Department, STFC Daresbury Laboratory, Warrington, WA4 4AD, United Kingdom.

## Abstract

*Salmonella* Typhimurium ST313 causes invasive nontyphoidal *Salmonella* (iNTS) disease in sub-Saharan Africa, targeting susceptible HIV^+^, malarial or malnourished individuals. An in-depth genomic comparison between the ST313 isolate D23580, and the well-characterized ST19 isolate 4/74 that causes gastroenteritis across the globe, revealed extensive synteny. To understand how the 856 nucleotide variations generated phenotypic differences, we devised a large-scale experimental approach that involved the global gene expression analysis of strains D23580 and 4/74 grown in sixteen infection-relevant growth conditions. Comparison of transcriptional patterns identified virulence and metabolic genes that were differentially expressed between D23580 versus 4/74, many of which were validated by proteomics. We also uncovered the *S.* Typhimurium D23580 and 4/74 genes that showed expression differences during infection of murine macrophages. Our comparative transcriptomic data are presented in a new enhanced version of the *Salmonella* expression compendium SalComD23580: bioinf.gen.tcd.ie/cgi-bin/salcom_v2.pl. We discovered that the ablation of melibiose utilization was caused by 3 independent SNP mutations in D23580 that are shared across ST313 lineage 2, suggesting that the ability to catabolise this carbon source has been negatively selected during ST313 evolution. The data revealed a novel plasmid maintenance system involving a plasmid-encoded CysS cysteinyl-tRNA synthetase, highlighting the power of large-scale comparative multi-condition analyses to pinpoint key phenotypic differences between bacterial pathovariants.

## Introduction

*Salmonella enterica* serovar Typhimurium (*S.* Typhimurium) infects a wide range of animal hosts, and generally causes self-limiting gastroenteritis in humans. Variants of this serovar, belonging to sequence-type ST313, are associated with invasive nontyphoidal *Salmonella* (iNTS) disease in susceptible HIV^+^, malaria-infected or malnourished individuals in sub-Saharan Africa [1]. iNTS causes around 681,000 deaths per year worldwide, killing 388,000 people in Africa alone [2]. The multidrug resistance of ST313 isolates complicates patient treatment and accounts for the high case fatality rate (20.6%) of iNTS disease [3]. Two ST313 lineages have been associated with iNTS, and the clonal replacement of lineage 1 by lineage 2 is hypothesized to have been driven by the gain of chloramphenicol resistance by lineage 2 [4]. Genetically-distinct ST313 isolates that do not belong to lineages 1 and 2 have been described in the United Kingdom [5] and in Brazil [6].

The globally-distributed *S.* Typhimurium sequence type ST19 causes gastroenteritis in humans and invasive disease in mice. Following oral ingestion, these bacteria colonise the gut and stimulate inflammation by a *Salmonella* pathogenicity island (SPI)-1-mediated process. Subsequently, ST19 can survive, and proliferate in a “*Salmonella*-containing vacuole” (SCV) within epithelial cells or macrophages that involves the SPI-2 type three secretion system responsible for systemic disease in mammalian hosts [7]. Host-restriction of other *Salmonella* pathovariants has been associated with genome degradation caused by pseudogene formation [8–11]. This process involves the loss or inactivation of virulence genes required for colonisation of the mammalian gut, whilst the ability to thrive inside macrophages is maintained.

Phenotypic differences between ST313 and ST19 have been summarized previously [12], and new studies have since been published. S1 Table lists 20 phenotypic features that differentiate ST313 from ST19 isolates, at the level of metabolism, motility, and stress resistance [13–24]. In terms of infection biology, reports of the relative ability of ST313 and ST19 isolates to invade epithelial cells and macrophages have yielded conflicting results (S1 Table) [6,13,15,17,25–27]. It is clear that ST313 infection of macrophages stimulates lower levels of cytotoxicity and inflammasome response than ST19 infections [13,25]. Following treatment with human serum, more complement was required for antibody-mediated bactericidal killing of ST19 than for ST313 isolates [14]. Animal infection experiments have demonstrated that ST313 isolates can infect non-human hosts, including mice, cows, chickens and macaques [15–18,28,29]. Taken together, these findings confirm that ST313 is a distinct pathovariant of *S.* Typhimurium [30]. However, the molecular mechanisms responsible for the phenotypic signature of ST313 pathovariant remain to be understood, and required a bespoke experimental approach.

D23580 is the ST313 lineage 2 reference strain, a typical representative Malawian isolate isolated from an HIV-negative child in 2004 [19]. We previously defined the transcription start sites (TSS) of this strain, and identified a SNP in the promoter of the *pgtE* gene specific to ST313 lineage 2 that modulated virulence [20]. To investigate whether the ability of the ST313 and ST19 sequence types of *S.* Typhimurium to cause different types of human disease was a genetic characteristic of the two types of bacteria, we identified all genomic differences between D23580 and 4/74. We then generated a comprehensive dataset for studying the mechanisms of infection-relevant differences between ST313 and ST19 listed in S1 Table. We hypothesised that transcriptional differences between the two strains would account for specific phenotypic differences and we present a multi-condition transcriptomic comparison of the ST313 strain, D23580, with the ST19 strain, 4/74 (S1 Fig).

## Results

### Resequencing and reannotation of D23580, the *Salmonella* Typhimurium ST313 reference strain

*S.* Typhimurium D23580 was the first ST313 isolate to be genome sequenced [19]. At that time, the presence of one D23580-specific plasmid, pBT1, was reported. To facilitate a robust transcriptomic analysis of D23580, we re-sequenced the strain using a combination of long-read PacBio and short-read Illumina technologies. Following a hybrid assembly approach (Materials and Methods), three contigs were identified: the 4,879,402 bp chromosome, the 117,046 bp pSLT-BT plasmid, and the 84,543 bp pBT1 plasmid (accession: XXXXXXXXXX). Comparison with the published D23580 genome (accession: FN424405) [19], identified just three nucleotide differences in the chromosome. Specifically, an extra nucleotide at the 304,327 position (1 bp downstream of Asp-tRNA), at the 857,583 position (1 bp upstream of Lys-tRNA), and one nucleotide change at position 75,492 (T-to-C; intergenic region) were identified. The sequence of the pSLT-BT plasmid had a single-nucleotide deletion difference at position 473, in an intergenic region. The sequence of the pBT1 plasmid has not been reported previously, and a primer-walking approach was used to sequence the two remaining small plasmids carried by D23580 (Materials and Methods), pBT2 and pBT3 (2556 bp and 1975 bp, respectively) (accession: XXXXXXXXXX).

To maximise the functional insights to be gained from a transcriptomic analysis, a well-annotated genome is required. The published annotation for D23580 dates back to 2009 [19], and lacked certain essential bacterial genes such as the two outer membrane proteins *lppA* and *lppB* [31]. Accordingly, we searched for important non-annotated bacterial genes, and used D23580 transcriptomic data (described below) to cross-reference the locations of transcripts with the location of coding genes (S1 Text). This analysis allowed us to update the published annotation of D23580 by adding 86 new coding genes, 287 sRNAs, and correcting the start or end locations of 13 coding genes (S2 Table). The re-sequenced and re-annotated *S.* Typhimurium D23580 genome is subsequently referred to as D23580_liv (accession: XXXXXXXXXX).

### The *S.* Typhimurium D23580 and 4/74 genomes are 95% identical

Previously the D23580 genome had been compared with the attenuated laboratory *S.* Typhimurium LT2 strain [19,32,33]. To assess the similarities and differences between the ST313 strain D23580 and a virulent ST19 isolate, a detailed comparative genomic analysis was performed against the ST19 strain 4/74 (S1 Text). 4/74 is a prototrophic *S.* Typhimurium ST19 strain that is highly virulent in four animal models [34], and is the parent of the widely-used SL1344 auxotrophic strain [35]. D23580 and 4/74 share 92% and 95% of coding genes and sRNAs, respectively (S2 Table). Genetic differences included 788 SNPs, 3 multi‐nucleotide polymorphisms (MNPs), 65 indels, as well as 77 D23580-specific pseudogenes that have been listed elsewhere [19]. Analysis of the SNPs, using the 4/74 annotation as a reference, showed that 379 were non-synonymous, 255 were synonymous, six were located in sRNAs, nine generated stop codons in coding genes and seven inactivated stop codons in D23580 (S3 Table). The final 132 SNPs were in intergenic regions. Fig 1 compares the chromosome and pSLT plasmid organisation of strains 4/74 and D23580, and shows the distribution of the indels and three SNP classes that differentiate the two strains. Seventeen of the SNPs and indels were located ≤ 40 nucleotides upstream of one of the D23580 TSS [20], raising the possibility of a direct influence upon the level of transcription.

**Fig. 1.**
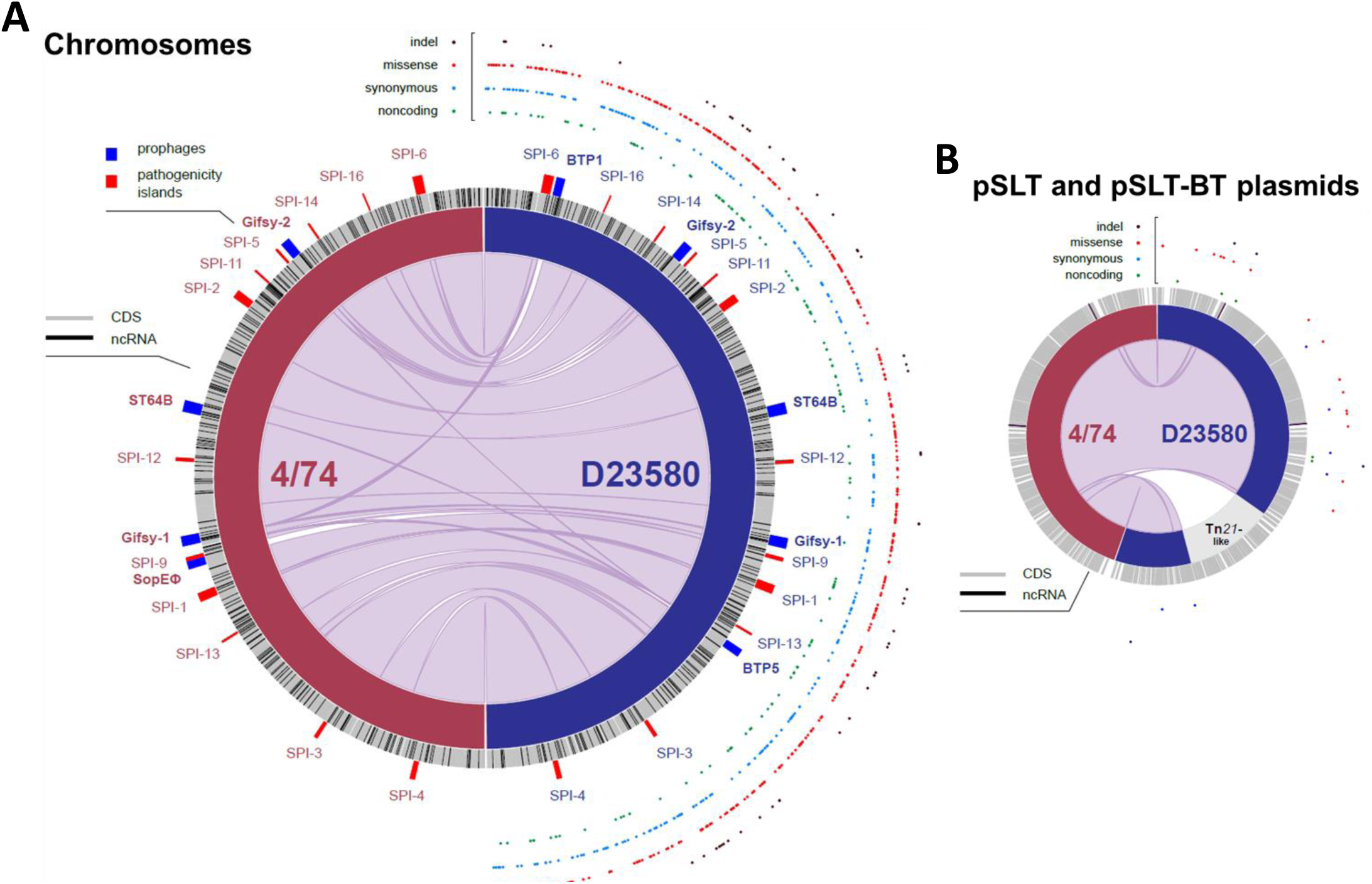
Comparative genomic analysis between *S.* Typhimurium 4/74 and D23580. Plots were obtained using the Circa software (http://omgenomics.com/circa/). (A) 4/74 and D23580 chromosomes; (B) 4/74 pSLT and D23580 pSLT-BT plasmids. In both panels, 4/74 data are represented on the left and D23580 data on the right. The four functional types of variants between D23580 and 4/74 are shown on the right hand side of each panel.

Regarding prophage complement, SopEϕ [36] was absent from D23580 and present in strain 4/74 [19,24]. As we established earlier, D23580 carries two ST313-specific prophages, BTP1 and BPT5 [19,24]. In terms of plasmids, the genome of 4/74 includes pSLT^4/74^, pCol1B9^4/74^, and pRSF1010^4/74^ [35]. In contrast, D23580 carries a distinct plasmid complement, namely pSLT-BT, pBT1, pBT2 and pBT3 [19]. The pSLT-BT plasmid of D23580 carries a Tn*21*-based insertion element that encodes resistance to five antibiotics [19].

The D23580 and 4/74 strains carry 4396 orthologous coding genes (S1 Text). Ten of the orthologues were encoded by the D23580-specific prophages BTP1 and BTP5, or by the 4/74-specific pRSF1010^4/74^ plasmid, and so were excluded from further analysis. A total of 279 orthologous sRNAs were found in both strains. The sRNA-associated differences included three 4/74-specific sRNAs (STnc3640, STnc1400, STnc3800), and the duplication of IsrB-1 in D23580. Eight new sRNAs were found in the BTP1 prophage region of D23580, and the existence of four was confirmed by northern blot (S2 Fig).

We identified 93 D23580-specific chromosomal genes that were encoded within prophage regions and absent from 4/74 (S2 Table): specifically, 59 BTP1 genes, 27 BTP5 genes, one Gifsy-2 gene and six Gifsy-1 genes. We found 89 4/74-specific chromosomal genes that were absent from D23580 (S2 Table). Most were located in the SopEϕ prophage region (68 genes), the Def2 remnant phage (13 genes), or three separate non-phage associated regions in D23580: *allB* (associated with allantoin utilization); the SPI-5 genes *orfX* and *SL1344_1032*; and an approximately 4 kb deletion that included genes *SL1344_1478* to *SL1344_82*.

A total of 4675 orthologous coding genes and non-coding sRNAs were shared by strains D23580 and 4/74. The sRNA IsrB-1 was removed as it was duplicated in D23580. To search for a distinct transcriptional signature of D23580, the expression levels of the 4674 orthologs was compared between D23580 and 4/74 using a transcriptomic approach.

### Comparison of transcriptional response to infection-relevant stress between *S.* Typhimurium ST313 D23580 and ST19 strain 4/74

To discover the similarities and differences in the transcriptome of strains D23580 and 4/74, we first used our established experimental strategy: the transcriptome of D23580 was determined using RNA isolated from 16 infection-relevant *in vitro* growth conditions [37], and during intra-macrophage infection [38,39]. To allow direct comparison of the D23580 transcriptomic data with strain 4/74, experiments were performed exactly as Kröger *et al*. (2013) and Srikumar *et al*. (2015) (Materials and Methods) [37,39].

The RNA-seq-derived sequence reads were mapped to the D23580_liv chromosome, and the pSLT-BT, pBT1, pBT2 and pBT3 plasmid sequences (Materials and Methods). Numbers of mapped sequence reads and other RNA-seq-derived statistical information are detailed in S4 Table. The level of expression of individual genes and sRNAs was calculated as transcripts per million (TPM) [40,41] for the chromosome, and the pSLT-BT and pBT1 plasmids (S5 Table). To achieve a complete transcriptomic comparison, we first re-analysed our 4/74 transcriptomic data [37,39] to add all transcripts expressed by the three plasmids, pSLT^4/74^, pCol1B9^4/74^, and pRSF1010^4/74^ (Materials and Methods, S4 Table).

Initial analysis focused on the expression characteristics of the strains D23580 and 4/74 in 17 distinct environmental conditions. The number of genes and sRNAs expressed in at least one condition for strain D23580 was 4365 (85%) out of 5110. 745 genes and sRNAs (15%) were not expressed in any of the 17 conditions. For strain 4/74, the number of genes and sRNAs that were expressed in at least one condition was 4306 (86%) out of 5026, consistent with our earlier findings [37] (S3 Fig). 3958 of the 4674 orthologous coding genes and sRNAs shared by strains D23580 and 4/74 were expressed in at least one growth condition in both strains.

A small minority (117) of orthologous genes were expressed in at least one condition in strain 4/74, but not in any of the conditions in D23580, with most showing low levels of expression (close to the threshold TPM = 10) (S5 Table). In contrast, we identified 82 orthologous coding genes and sRNAs that were expressed in at least one of the 17 growth conditions for D23580, but not expressed in 4/74 (S5 Table).

To compare the expression profiles of D23580 and 4/74, we made 17 individual pair-wise comparisons between the 17 growth conditions with the two strains (Materials and Methods, S3 Fig). The data confirmed that *S.* Typhimurium reacts to particular infection-relevant stresses with a series of defined transcriptional programmes that we detailed previously [37]. By comparing the transcriptomic response of two pathovariants of *S.* Typhimurium, the conservation of the transcriptional response is apparent (S3 Fig).

A complementary analytical approach was used to identify the transcriptional differences that relate to the distinct phenotypes of the ST313 and ST19 pathovariants (S1 Table). Overall, 1031 of the orthologous coding genes and sRNAs were differentially expressed (≥ 3 fold-change) between strains D23580 and 4/74, in at least one growth condition (Fig 2A). Transcriptional differences are highlighted in S3 Fig.

**Fig. 2.**
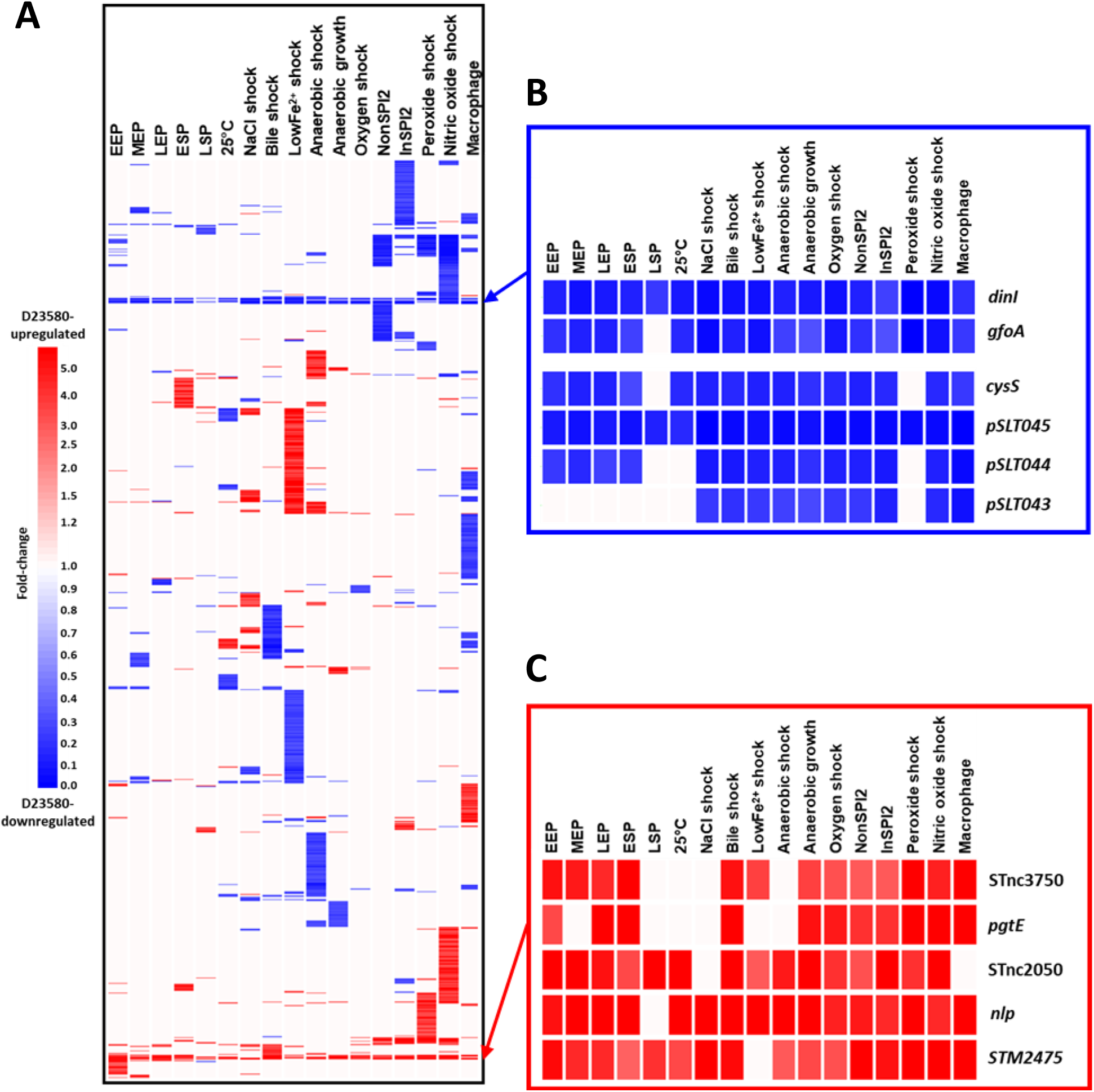
Inter-strain transcriptomic comparison of *S.* Typhimurium D23580 versus 4/74. Expression of orthologous coding genes and sRNAs was compared between strains D23580 and 4/74 during growth in 17 infection-relevant conditions. The transcriptional expression value (TPM) for each coding gene and sRNA in each condition in D23580 was divided by the TPM value for the same gene/sRNA and condition in 4/74. Heat maps were obtained using the GeneSpring GX7.3 software (Agilent). Cluster analysis was performed using data with ≥ 3 fold-change. (A) Heat map of the 1031 coding genes and sRNAs that showed significant difference (≥ 3 fold-change) between the two strains in at least one condition. (B) Heat map representing D23580-downregulated genes observed in all or most growth conditions. (C) Heat map of the D23580-upregulated coding genes and sRNAs observed in most growth conditions.

The terms “D23580-upregulated” and “D23580-downregulated” refer to genes that are either more or less expressed in D23580, compared to 4/74. Three coding genes were D23580-upregulated and six genes were D23580-downregulated in almost all growth conditions (Fig 2B and 2C). The up-regulated genes included *pgtE*, a gene that is highly expressed in D23580, responsible for resistance to human serum killing and linked to virulence [20]. The other two upregulated genes were *nlp*, encoding a *ner*-like regulatory protein, and the *STM2475 (SL1344_2438*) gene which encodes a hypothetical protein.

Three of the genes that were D23580-downregulated in most conditions (*pSLT043-5*) were located downstream of the Tn*21*-like element in the pSLT-BT plasmid (S4 Fig). Because the Tn*21*-like multidrug resistance island was inserted between the *mig-5* promoter region and the *pSLT043-5* genes, we hypothesise that the differential expression reflects transcriptional termination mediated by the Tn*21* cassette. Two other D23580-downregulated genes were located in the Gifsy-1 prophage region, *dinI-gfoA*. The presence of a SNP in the promoter D23580 P*_dinI-gfoA_* has already been proven to be responsible for the lack of viability of the Gifsy-1 phage in D23580 [24]. The final gene that was D23580-downregulated in most growth conditions was the *cysS* chromosomal gene, which encodes a cysteinyl-tRNA synthetase. Aminoacyl-tRNA synthetases are generally essential genes, required for cell growth and survival [42]. The unexpected low level of *cysS* expression in D23580 in several growth conditions (TPM values ranging from 5 to 18 excluding the late stationary phase and shock conditions) was investigated further (see below).

Intriguing patterns of differential expression were observed between strains D23580 and 4/74 in particular growth conditions for certain functional groups of *Salmonella* genes. For example, the flagellar regulon and associated genes showed a characteristic pattern of expression in the phosphate carbon nitrogen (PCN)-related minimal media and inside macrophages (S5 Fig). To allow us to make statistically significant findings, a larger-scale experiment was designed.

### Identification of the transcriptional signature of *S.* Typhimurium ST313 D23580

To generate a robust transcriptional signature of D23580, we focused on the five environmental conditions with particular relevance to *Salmonella* virulence, namely ESP (early stationary phase), anaerobic growth, NonSPI2 (SPI2-non-inducing), InSPI2 (SPI2-inducing) conditions and intra-macrophage. The ESP and anaerobic growth conditions stimulate expression of the SPI-1 virulence system, and SPI-2 expression is induced by the InSPI2 and macrophage conditions [37,39]. RNA was isolated from three biological replicates of both D23580 and 4/74 grown in the four *in vitro* environmental conditions. The three biological replicates were generated in parallel, in a new set of experiments. Additionally, RNA was extracted from two additional biological replicates of intra-macrophage *S.* Typhimurium, following infection of murine RAW264.7 macrophages for both D23580 and 4/74. Following RNA-seq, the sequence reads were mapped to the D23580 and 4/74 genomes using our bespoke software pipeline (Materials and Methods). The RNA-seq mapping statistics are detailed in S4 Table. To ensure that biologically meaningful gene expression differences were reported, we used very conservative cut-offs to define differential expression (Materials and Methods).

Following RNA-seq analysis of the three biological replicates of D23580 and 4/74 in five growth conditions, differential expression analysis of orthologous genes and sRNAs was performed with a rigorous statistical approach (Materials and Methods, S6 Table). We identified 677 genes and sRNAs that showed ≥ 2 fold-change (FDR ≤ 0.001) in at least one growth condition (Fig 3A). Between 6% (anaerobic growth) and 2% (InSPI2 condition) of orthologous genes and sRNAs were differentially expressed between the two strains (Fig 3A and 3B).

**Fig. 3.**
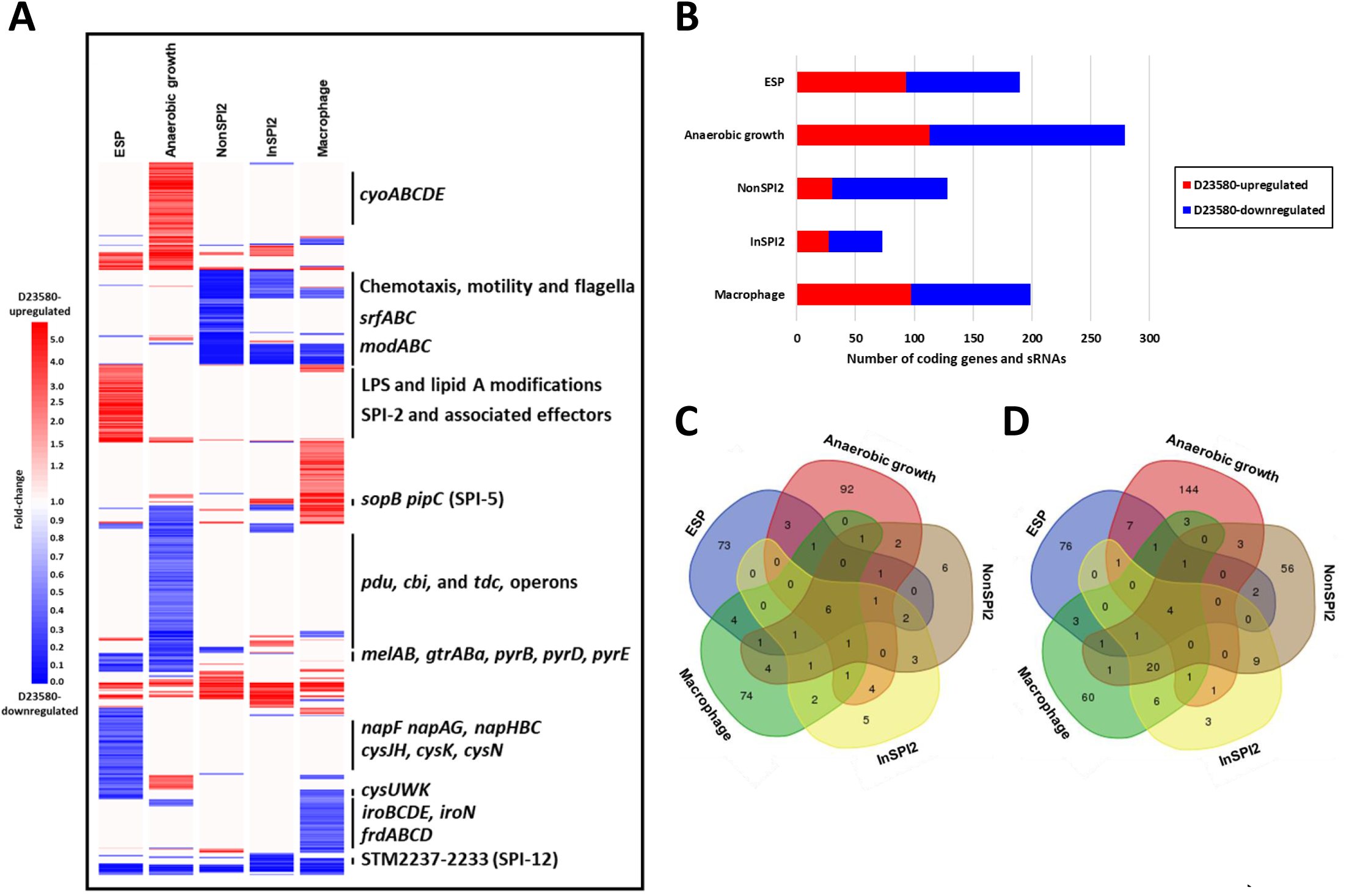
Transcriptional signature of *S.* Typhimurium D23580. Differential expression analysis of orthologous coding genes and sRNAs between strains D23580 and 4/74 during growth in five infection-relevant conditions. (A) Heat map highlighting biological relevant clusters. The CPM values of three biological replicates for each coding gene and sRNA in each condition in D23580 were compared to the CPM values for the same gene/sRNA and condition in 4/74. The heat map was obtained using GeneSpring GX7.3 (Agilent). Cluster analysis was performed with CPM values of the 677 coding genes and sRNAs that showed ≥ 2 fold-change and ≤ 0.001 of FDR in the differential expression analysis generated using Degust in at least one condition. (B) Number of coding genes and sRNAs differentially expressed in each of the five growth conditions, based on Degust results (≥ 2 fold-change, ≤ 0.001 FDR). (C) Venn diagram of the D23580-upregulated genes in the five growth conditions (http://bioinformatics.psb.ugent.be/webtools/Venn/). (D) Venn diagram of the D23580-downregulated genes in the five growth conditions.

The ability to swim in semi-solid agar is a key phenotypic difference between D23580 and 4/74 [13,16]. We confirmed that D23580 was less motile than 4/74 (S5 Fig), but did not observe significant differences in motility gene expression in complex media between the strains at the transcriptional level (S5 Fig). One nucleotide deletion and 11 SNP differences were found in the flagellar regulon between the two strains: one mutation in the promoter region of *mcpA;* three synonymous mutations in *flgK, cheA*, and *fliP;* four non-synonymous mutations in *flhA, flhB, fliB*, and *mcpC;* and three mutations in the 5’UTRs of *motA, flhD*, and *mcpA*. FlhA and FlhB are transmembrane proteins that are essential for flagellar protein export [43]. The SNP in *flhB* is 4/74-specific, as other ST19 strains, such as LT2 and 14028, conserved the SNP in D23580. The D23580 *flhA* SNP was specific to ST313 lineage 2.

To investigate the function of the 4/74 *flhA* SNP, the mutation was introduced to the chromosome of D23580 by single nucleotide engineering (D23580 *flhA^4/74^*). Motility of the D23580 *flhA^4/74^* mutant was significantly increased compared to the D23580 wild-type strain (S5 Fig). We originally hypothesized that the *flhA* SNP was related to the reported decreased inflammasome activation in macrophages, which is thought to contribute to the stealth phenotype of *S.* Typhimurium ST313 that involves evasion of the host immune system during infection [25]. However, no significant differences in cell death due to inflammasome activation were found between D23580 wild type and the D23580 *flhA^4/74^* mutant (S5 Fig).

The transcriptomic data did offer an explanation for the reduced motility of D23580 on minimal media. In the NonSPI2 condition, all flagellar genes were D23580-downregulated, with the exception of the master regulators *flhDC* (S5 Fig). In contrast, in the InSPI2 and intra-macrophage condition, only the flagellar class 2 genes (such as *flgA*) were significantly down-regulated. RflP (YdiV) is a post-transcriptional negative-regulator of the flagellar master transcriptional activator complex FlhD_4_C_2_ [44–46]. We speculate that the downregulation of the flagellar regulon in NonSPI2 could be due to a significant upregulation (3.5 fold-change) of *rflP* in this low-nutrient environmental condition. This differential expression was not seen in the InSPI2 growth condition which only differs from NonSPI2 by a lower pH (5.8 versus 7.4) and a reduced level of phosphate [37].

We identified six genes and sRNAs that were D23580-upregulated in most growth conditions, specifically *pgtE, nlp, ydiM, STM2475 (SL1344_2438*), the ST64B prophage-encoded *SL1344_1966*, and the sRNA STnc3750 (Fig 3C). Just four genes were D23580-downregulated in all conditions, namely *pSLT043-5 (SLP1_0062-4*), and *cysS* (Fig 3D). These findings confirmed that biologically significant information can be extracted from the initial 17-condition experiment (Fig 2B and 2C) as similar genes were up/down-regulated across the multiple conditions of the replicated experiment.

The transcriptomic data were interrogated to identify virulence-associated genes that were differentially expressed between D23580 and 4/74. Coding genes and sRNAs located within the *Salmonella* pathogenicity islands SPI-1, SPI-2, SPI-5, SPI-12 and SPI-16 showed differential expression between D23580 and 4/74, in at least one growth condition (S6 Fig). The SPI-5-encoded *sopB* gene (encoding a SPI-1 effector protein) and its associated chaperone gene (*pipC*) were significantly D23580-upregulated in the InSPI2 and intra-macrophage conditions. In contrast, the SPI-12-associated genes *STM2233-7 (SL1344_2209-13*) were D23580-downregulated in the same two growth conditions. Most SPI-2 genes were significantly D23580-upregulated in the ESP condition, raising the possibility that the non-induced level of expression of SPI2 is higher in D23580 than 4/74.

The most highly differentially expressed genes in ESP (≥ 4 fold-change, FDR ≤ 0.001) (S6 Fig), included the D23580-upregulated genes required for itaconate degradation (*ripC), myo*-inositol utilization (*reiD*), and proline uptake (*putA*). D23580-downregulated genes in the same growth condition included those involved in uptake of uracil and cytosine (*uraA* and *codB*), melibiose utilization (*melAB*), carbamoyl phosphate metabolism and pyrimidine biosynthesis (*carAB* and *pyrEIB*), nitrate reductase (*napDF*), and sulfate metabolism (*cysPU* and *sbp*).

Genes that were differentially expressed between D23580 and 4/74 were also identified during infection of RAW264.7 macrophages. The 16 genes that were most highly D23580-upregulated (≥ 4 fold-change, FDR ≤ 0.001) included a β-glucosidase, *STM3775 (SL1344_3740);* genes involved in cysteine metabolism, *cdsH;* oxidation of L-lactate, *STM1621 (SL1344_1551);* and the transcriptional regulator *rcsA*. Genes that were D23580-downregulated during infection of macrophages were involved in the secretion and import of siderophores (*iroC* and *iroD*), uptake of sialic acid (*nanM*), and maltose or maltodextrin (*malEFK*), and *STM1630 (SL1344_1560*), which encodes a hypothetical protein.

The key features of the transcriptional signature of D23580 included the differential expression of the flagellar and associated genes, genes involved in aerobic and anaerobic metabolism, and iron-uptake genes. Specifically, the aerobic respiratory pathway *cyoABCDE* was D23580-upregulated in the anaerobic growth condition, and anaerobic-associated pathways *pdu, cbi* and *tdc* operons were D23580-downregulated. Importantly, genes associated with the acquisition of iron through production and uptake of siderophores were D23580-downregulated in the intra-macrophage environment. In summary, the transcriptional signature of D23580 suggests that the biology of ST313 lineage 2 differs from ST19 under anaerobic conditions *in vitro* and during infection of murine macrophages.

The challenge of data reproducibility in experimental science is widely acknowledged [47,48]. To assess the robustness of our experiments, the RNA-seq-derived expression profiles that we generated from five replicated conditions were compared with five relevant individual conditions. There was a high level of correlation between the individual versus replicated datasets (correlation coefficients between 0.88 and 0.97) (S7 Fig). However, different levels of expression were seen between the individual and the replicated ESP growth condition of D23580 for a small minority of genes. The main variations in terms of functional gene groups involved cysteine metabolism, carbamoyl-phosphate and pyrimidine biosynthesis, and nitrate metabolism. Variation in expression of the *ripCBA-lgl* operon was also observed during anaerobic growth. We speculate that these alterations in gene expression reflect experimental variations such as the use of different batches of media.

### 1.5% of proteins were differentially expressed between D23580 and 4/74

RNA-seq-based transcriptomic analysis does not reflect the translational and post-translational levels of regulation [49]. To identify proteins that differentiated strains D23580 and 4/74, we used a proteomic strategy that involved an LC-MS/MS platform, and analysed proteins from D23580 and 4/74 bacteria grown in the ESP condition (S7 Table). A label-free quantification approach identified 66 differentially expressed orthologous proteins (≥ 2 unique peptides, ≥ 2 fold-change, *p*-value < 0.05) (Fig 4), including 54 D23580-upregulated proteins and 12 D23580-downregulated proteins. The most highly D23580-upregulated protein was PgtE, corroborating our previous study [20]. Up-regulated proteins included those required for carbamoyl-phosphate and pyrimidine biosynthesis (CarAB and PyrIB), some SPI-1 proteins and associated effectors (PrgH, SipAB, InvG, SlrP, SopB, SopE2, SopA, SopD), RipAC (itaconate degradation) and Lgl (methylglyoxal detoxification).

**Fig. 4.**
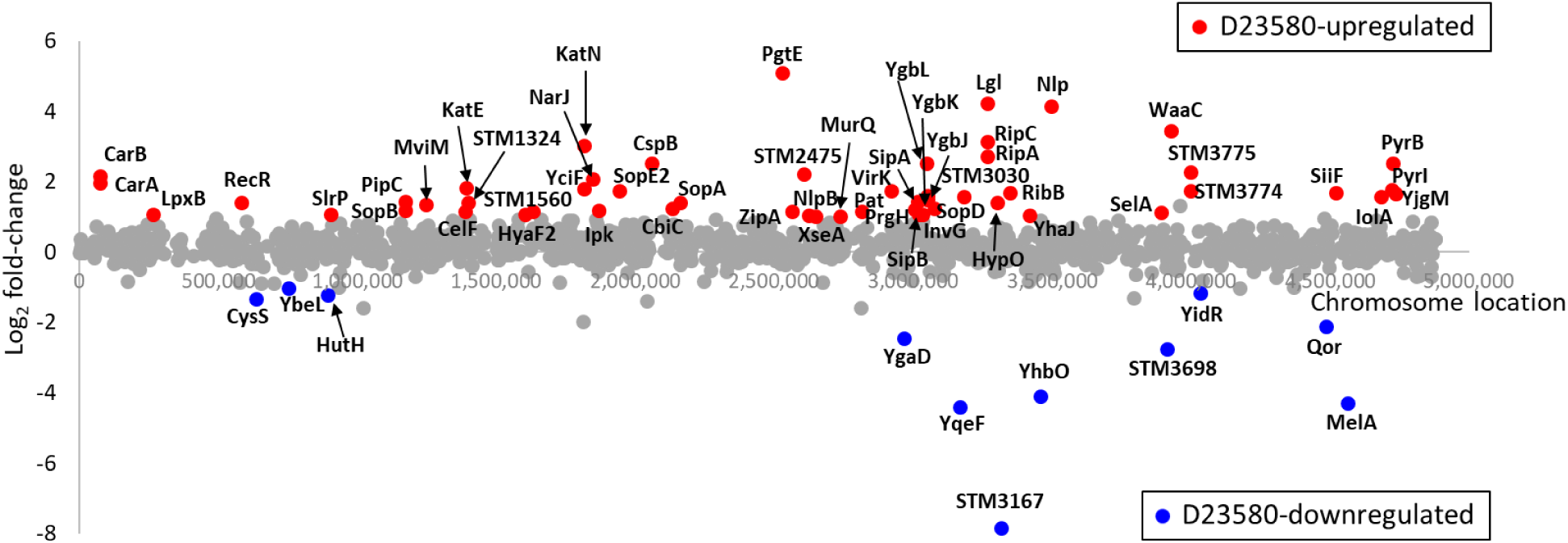
Differentially expressed proteins between *S.* Typhimurium D23580 and 4/74. Representation of significant D23580-upregulated proteins (red dots) and D23580-downregulated proteins (blue dots) in the ESP growth condition by Log_2_[fold-change] and the chromosome location in D23580 (≥ 2 unique peptides, ≥ 2 fold-change, *p*-value < 0.05). Grey dots refer to proteins that show non-significant differences.

To identify genes that were differentially expressed at both the transcriptional and translational levels, the quantitative proteomic data were integrated with the transcriptomic data. Eight D23580-upregulated proteins (YciF, SopA, PgtE, STM2475, RipC, RibB, Nlp, and STM3775) were significantly up-regulated in the transcriptomic data (≥ 2 fold-change, FDR ≤ 0.001). Four differentially expressed proteins (pSLT043, CysS, YgaD, MelA) were D23580-downregulated at the transcriptomic level (≥ 2 fold-change, FDR ≤ 0.001) (Fig 5). Overall, 12 genes were differentially expressed at both the transcriptional and protein levels.

**Fig. 5.**
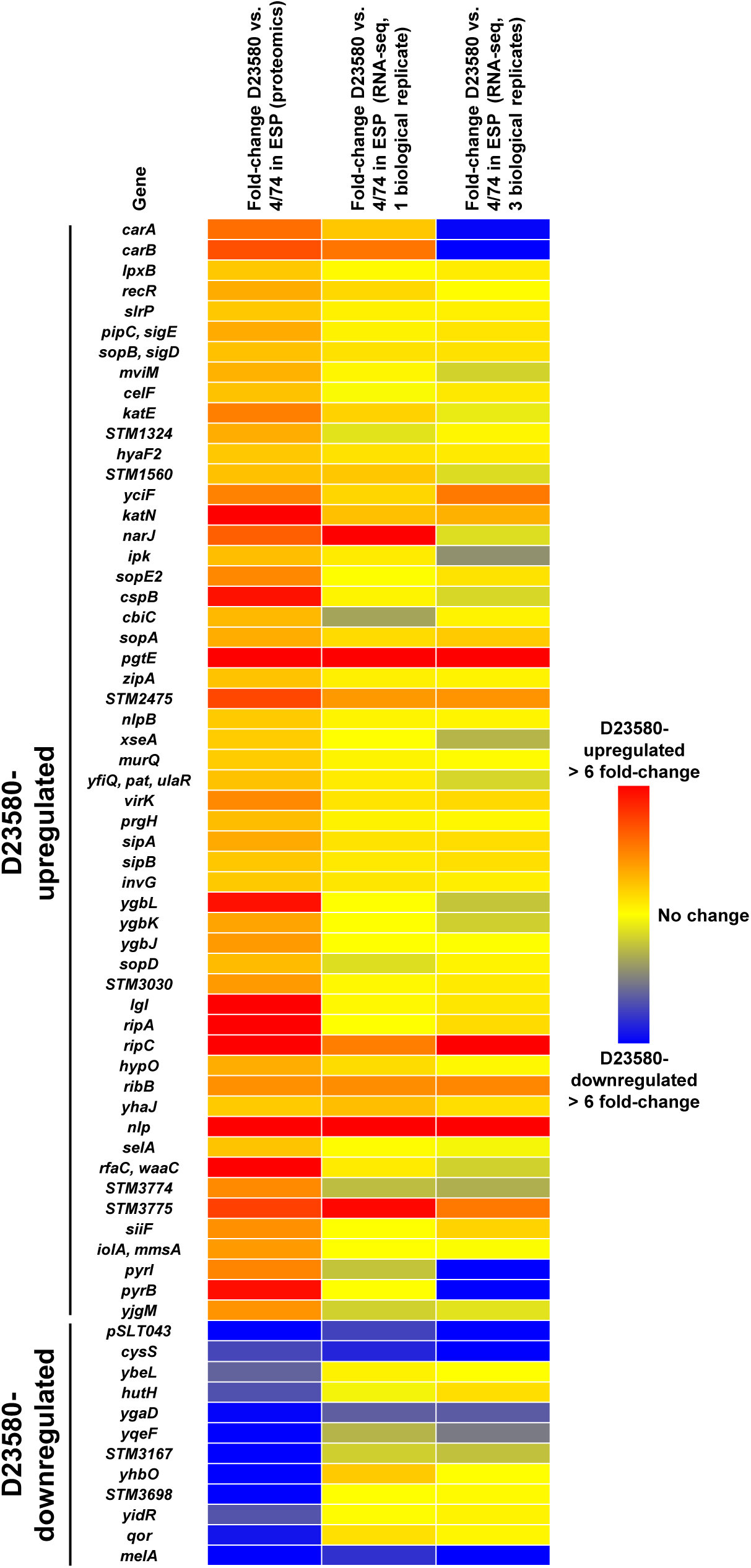
Heat map of the 66 differentially expressed proteins between *S.* Typhimurium 4/74 and D23580. Data represent levels of expression at the proteomic level in the ESP growth condition, and in two independent ESP RNA-seq datasets (one biological replicate versus three biological replicates).

### Evolution of *S.* Typhimurium ST313 involved the SNP-based inactivation of melibiose utilization genes

The melibiose utilization system consists of three genes: *melR*, which encodes an AraC-family transcriptional regulator; *melA*, encoding the alpha-galactosidase enzyme; and *melB*. MelB is responsible for the active transport of melibiose across the bacterial cell membrane. We found that the *melAB* genes were D23580-downregulated at the transcriptomic level (Fig 6A). The differential expression of *melA* was confirmed at the proteomic level (Fig 4A).

**Fig. 6.**
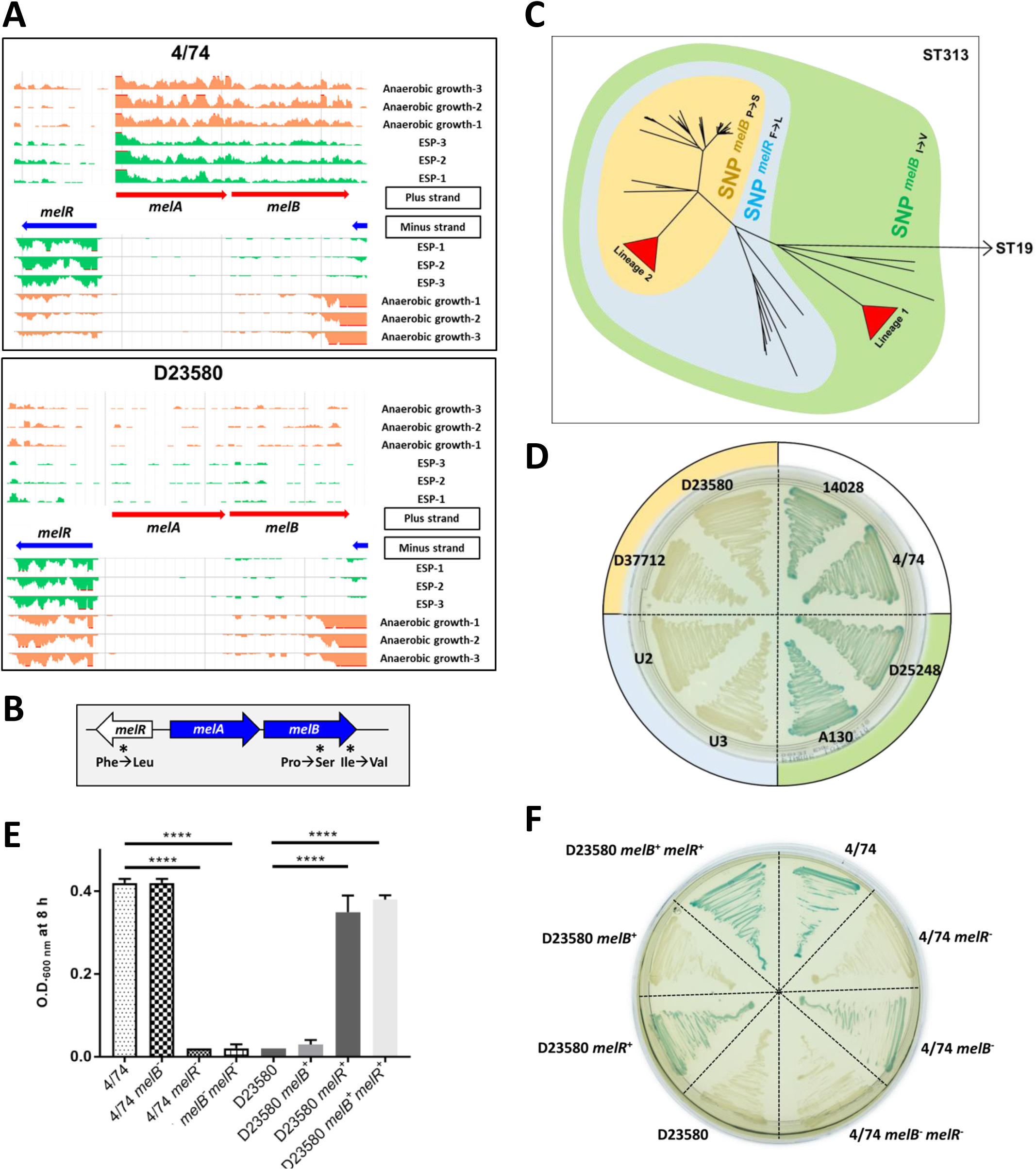
Melibiose phenotype differentiates the *S.* Typhimurium ST19 and ST313 strains. (A) Visualization of RNA-seq data with three biological replicates in the ESP and anaerobic growth conditions using JBrowse [50] for the melibiose utilization operon. Scale of the mapped reads was 1 to 100. (B) Presence of three non-synonymous SNPs in the melibiose utilization genes (4/74 → D23580). (C) Accumulation of SNPs in the melibiose utilization genes during the evolution of ST313, in context of a whole genome core SNP phylogeny. Isolate names, ST313 lineage and genotype for the three SNPs are included in S8 Table. (D) Alpha-galactosidase activity of representatives of ST19 and ST313 strains on Pinnacle *Salmonella* ABC medium (LabM), green = positive, colorless = negative. The colors of the external circle correlate with the colors represented in the tree in “C”. (E) Bacterial growth in minimal M9 medium supplemented with 0.4% melibiose. Bars represent the mean of seven biological replicates and standard deviation. Significant differences (****) indicate *p*-value < 0.0001. (F) Alpha-galactosidase activity of 4/74 and D23580 wild-type strains and corresponding mutants. The ability to use melibiose is rescued in D23580 by exchange of the three SNP mutations.

In strain D23580, the melibiose utilization genes contain three non-synonymous SNPs (4/74 → D23580). Two are present in *melB* (Pro → Ser at the 398 AA, Ile → Val at the 466 AA) and one in *melR* (Phe → Leu) (Fig 6B). The three SNPs were analysed in the context of a phylogeny of 258 genomes of *S.* Typhimurium ST313 that included isolates from Malawi, as well as more distantly-related ST313 genomes from the UK [5] (S8 Table). All three SNPs were found to be monophyletic, allowing us to infer the temporal order in which they arose and representing an accumulation of SNPs in melibiose utilization genes over evolutionary time. The first SNP, *melB* I466V, was present in all 258 ST313 strains tested and therefore arose first. The second SNP, in *melR*, was present in all ST313 lineage 2 and UK-ST313 genomes, suggesting that it appeared prior to the divergence of these phylogenetic groups [5]. The final SNP, *melB* P398S, is present in all ST313 lineage 2 and a subset of UK-ST313 genomes, consistent with this being last of the three mutations to arise (Fig 6C). ST313 strains can therefore be classified into groups of strains containing one, two or three SNPs in melibiose utilization genes.

It has been reported that D23580 did not ferment melibiose whereas a ST313 lineage 1 isolate (A130), *S.* Typhimurium SL1344 and *S.* Typhi Ty2 were able to utilize melibiose as a sole carbon source [18]. MelB catalyzes the symport of melibiose with Na^+^, Li^+^, or H^+^ [51]. We confirmed that ST19 strains, and strains belonging to the ST313 lineage 1, were positive for alpha-galactosidase activity. In contrast, isolates representing the UK-ST313 lineage and the ST313 lineage 2 were unable to utilize melibiose.

To determine the biological role of the SNPs in the *melB* and *melR* genes, we employed a genetic approach. Single nucleotide engineering was used to generate isogenic strains that reflect all three melibiose gene SNP states for determination of the role of the SNP differences between ST313 lineage 2 and ST19 in the alpha-galactosidase (MelA)-mediated phenotypic defect (Fig 6D). Melibiose utilization in D23580 was rescued by nucleotide exchange of the three SNP mutations (D23580 *melB*^+*melR*+^) (Fig 6E). D23580 recovered its ability to grow with melibiose as the sole carbon source after exchanging only the *melR* SNP with 4/74 (D23580 *melR*^+^). In contrast, D23580 did not grow in the same medium when the exchange only involved the two *melB* SNPs (D23580 *melB*^+^). 4/74 lost its ability to utilize melibiose as sole carbon source when we introduced the D23580 *melR* SNP (4/74 *melR*^-^ and 4/74 *melB*^-^ *melR*^-^). However, an exchange of the two nucleotides in *melB* did not eliminate the ability of 4/74 to grow in minimal medium with melibiose (4/74 *melB*^-^). These data correlated with the alpha-galactosidase activity of the mutants, although a slight difference was observed between strains D23580 *melR*^+^ (light green) and D23580 *melB*^+^ *melR*^+^ (green), and between strains 4/74 (green) and 4/74 *melB*^-^ (light green) (Fig 6F) suggesting an altered efficiency of melibiose utilization between the two strains. To completely restore alpha-galactosidase activity in D23580, the reversion of the non-synonymous SNPs in both the *melR* and *melB* genes was required. Our data suggest that the *melR* SNP is critical for the loss of function of the melibiose utilization system.

In a chicken infection model, the *S.* Typhimurium *melA* transcript is more highly expressed in the caecum than during *in vitro* growth [52]. In a chronic infection model, accumulation of melibiose was observed in the murine gut after infection with *S.* Typhimurium [53]. The combination of the SNP-based inactivation of melibiose catabolism with the conservation of the key SNPs in ST313 lineage 2 is consistent with a functional role in ST313 virulence, and we are currently examining this possibility.

### A plasmid-encoded cysteinyl-tRNA synthetase is required for growth in D23580

The dramatic down-regulation of the chromosomal *cysS* gene at both the transcriptomic (Fig 7A) and proteomic levels (Fig 4A) was studied experimentally. The coding and non-coding regulatory regions of the chromosomal *cysS* were identical at the DNA level in strains D23580 and 4/74. The chromosomal *cysS* gene encodes a cysteinyl-tRNA synthetase, which is essential for cell growth in *S.* Typhimurium and other bacteria [42,54,55]. To investigate *cysS* gene function, we consulted a transposon-insertion sequencing (TIS) dataset for *S.* Typhimurium D23580 (manuscript in preparation). Genes that show the absence or low numbers of transposon insertion sites are considered to be ‘required’ for bacterial growth in a particular condition [55,56]. The data suggested that a functional chromosomal *cysS* was not required for growth in rich medium (Fig 7B). We searched for *S.* Typhimurium D23580 genes that encoded a cysteinyl-tRNA synthetase, and identified the pBT1-encoded gene, *pBT1-0241 (cysS^pBT1^*), which the TIS data suggested to be ‘required’ for growth in rich medium (Fig 7B).

**Fig. 7.**
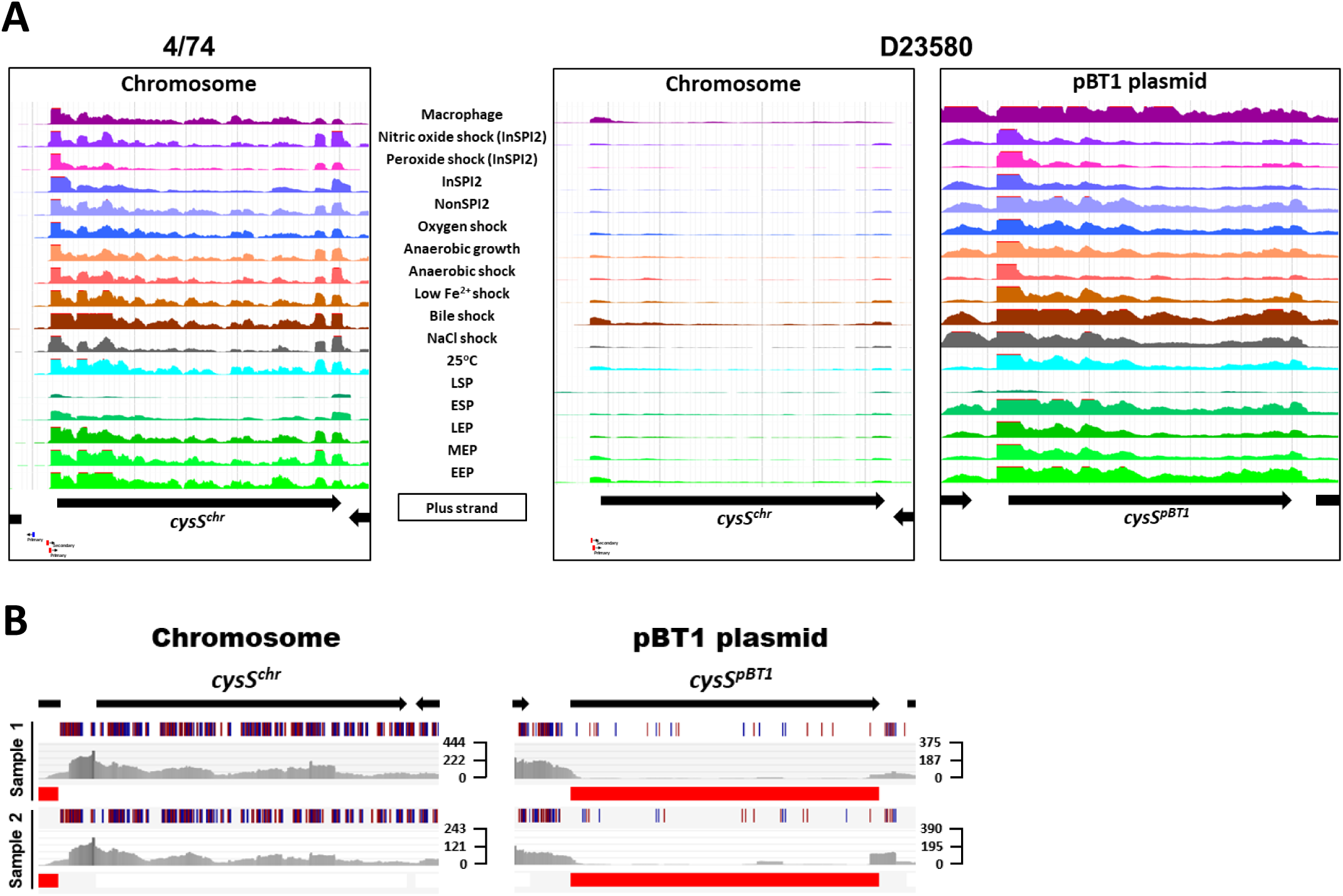
The pBT1 plasmid encodes the functional cysteinyl-tRNA synthetase in *S.* Typhimurium D23580. (A) RNA-seq data for *cysS* in 4/74, and chromosomal and pBT1-plasmid-encoded *cysS* in D23580 from the online JBrowse resources provided in this study. Scale of the mapped reads was 1 to 500. (B) Transposon library results for the *cysS^chr^* and *cysS^pBT1^* genes in D23580. Figures were obtained using the Dalliance genome viewer [57]. Black arrows at the top represent genes. Each sample is represented by three tracks. The first track contains blue and red lines that correspond to transposon insertion sites; blue means = orientation of the transposon is the same as the direction of the gene, red = opposite direction. The second track shows raw data for the Illumina sequencing reads. The third track highlights in red those genes that were considered ‘required’ for growth in that condition based on an insertion index. The insertion index was calculated for each gene as explained in [56,58], and genes with insertion index values < 0.05 were considered as ‘required’ for growth in the Lennox rich medium. The scale on the right represents sequence read coverage.

To investigate cysteinyl-tRNA synthetase function in D23580, individual knock-out mutants were constructed in the chromosomal *cysS* gene (*cysS^chr^*), and the *cysS^pBT1^* gene. These genes were 89% identical at the amino acid level and 79% at the nucleotide level. The *cysS^pBT1^* mutant was whole genome sequenced to confirm the absence of secondary unintended mutations. The pBT1 plasmid was also cured from D23580. We determined the relative fitness of the two *cysS* mutants and the pBT1-cured strain. The D23580 wild type, D23580 *ΔcysS^chr^* and D23580 ΔpBT1 mutants grew at similar rates in LB, whilst the D23580 *ΔcysSp^BT1^* mutant showed an extended lag phase (Fig 8A). The D23580 *ΔcysSp^BT1^* mutant showed a more dramatic growth defect in minimal medium with glucose as the sole carbon source (Fig 8B).

**Fig. 8.**
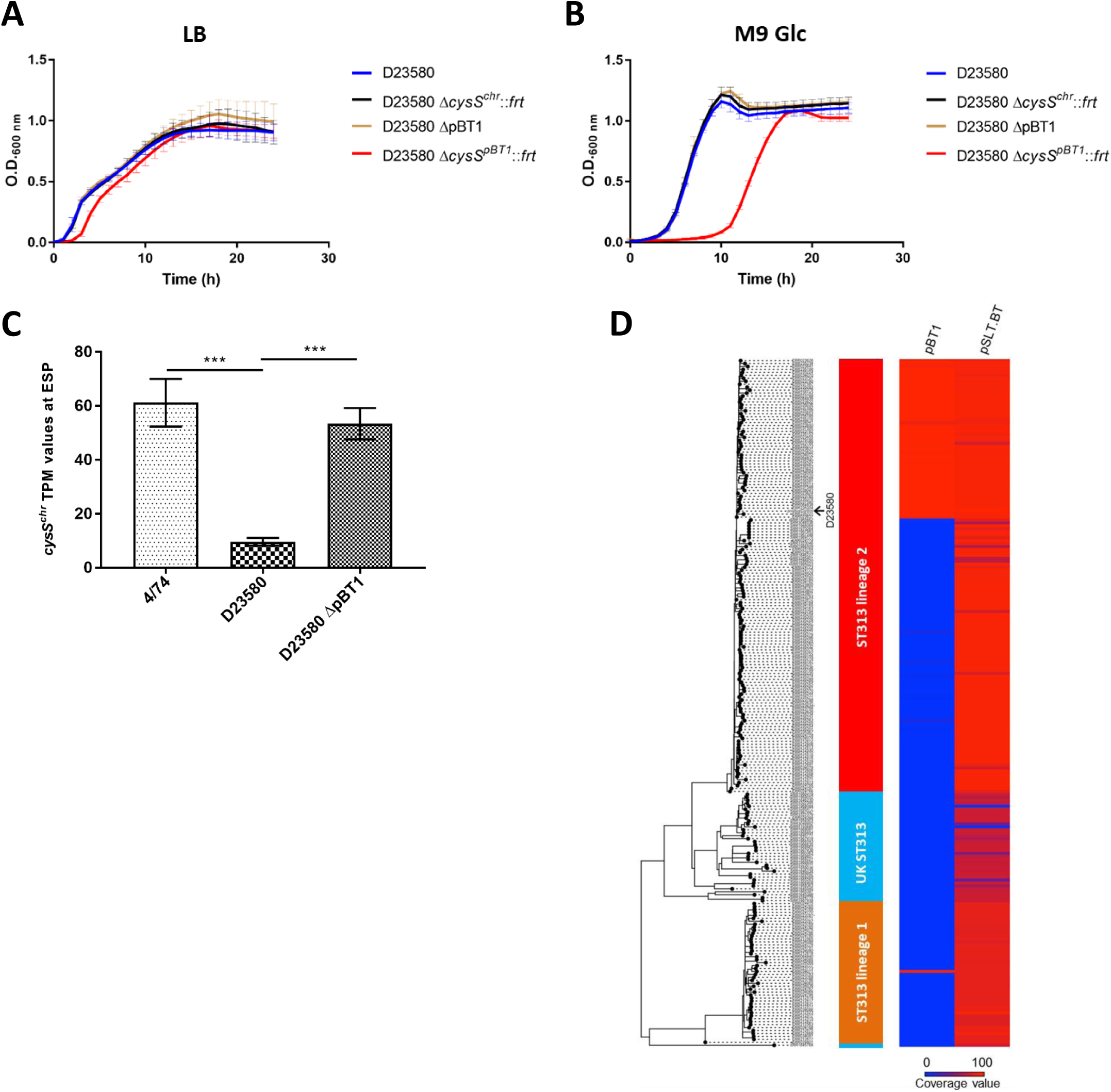
The pBT1-encoded *cysS* is required for optimal growth of *S.* Typhimurium D23580. (A) Growth curves of D23580 WT, D23580 Δ*cysS^chr^*::*frt*, D23580 pBT1-cured strain, and D23580 Δ*cysS^pBT1^*::*frt* strains in LB medium, n = 8 (standard deviations are represented). (B) Growth curves in minimal M9 medium supplemented with 0.4% glucose, n = 5 (standard deviations are represented). (C) Comparison of *cysS^chr^* expression levels (TPM values) of 4/74 (n = 3), D23580 (n = 3), and the D23580 pBT1-cured strain (n = 2) in the ESP growth condition. Bars represent mean values and standard deviations. Significant differences (***) indicate *p*-value < 0.001. (D) The pBT1 plasmid is present in a subset of ST313 isolates of lineage 2 and in one isolate from lineage 1. Isolate names, ST313 lineage and coverage value for pBT1 and pSLT-BT are included in S8 Table.

To determine whether the presence of the pBT1 plasmid was linked to the decrease in *cysS^chr^* expression, RNA from two biological replicates was isolated from the pBT1-cured strain in the ESP growth condition. Differential expression analysis between this mutant and the wild-type D23580 strain showed a significant increase in expression of *cysS^chr^*, with TPM levels close to those seen in 4/74 (Fig 8C, S5 Table). These results suggested the pBT1 plasmid is responsible for the down-regulation of *cysS^chr^* expression in D23580.

The conservation of pBT1 was studied among 233 ST313 strains and compared to the presence of the pSLT-BT plasmid which was found in all lineage 2 isolates (Fig 8D, S8 Table). Approximately 37% of ST313 lineage 2 isolates carried the pBT1 plasmid. The pBT1 plasmid has rarely been seen previously, but did show significant sequence similarity to five plasmids found in *Salmonella* strains isolated from reptiles and elsewhere (98 to 99% nucleotide identity over 92 to 97% of the plasmid sequence, accessions JQ418537, JQ418539, CP022141, CP022036 and CP022136, S1 Text).

Examples of essential bacterial genes located on plasmids are rare and this phenomenon has been previously explored [59]. We conclude that the essentiality of the *cysS^pBT1^* gene provides a novel strategy for plasmid maintenance in a bacterial population.

### The SalComD23580 community data resource

To allow scientists to gain new biological insights from analysis of this rich transcriptomic dataset, we have made it available as an online resource for the visualization of similarities and differences in gene expression between ST313 (D23580) and ST19 (4/74), using an intuitive heat-map-based approach (bioinf.gen.tcd.ie/cgi-bin/salcom_v2.pl). To examine the transcriptional data in a genomic context, we generated two strain-specific online browsers that can be accessed from the previous link: one for D23580 and one for 4/74. The value of this type of online resource for the intuitive interrogation of transcriptomic data has been described recently [60].

## Perspective

To investigate the functional genomics of *S.* Typhimurium ST313, we first re-sequenced and re-annotated the genome of the D23580 isolate. Our comparative genomic analysis of two *S.* Typhimurium ST313 and ST19 isolates confirmed the findings of Kingsley *et al.* (2009) [19], identifying 856 SNPs and indels, many instances of genome degradation and the presence of specific prophages and plasmids. To discover the genetic differences that impact upon the biology of *S.* Typhimurium ST313, we used a functional transcriptomic approach to show that the two *S.* Typhimurium pathovariants shared many responses to environmental stress.

By investigating global gene expression in multiple infection-relevant growth conditions, we discovered that 677 genes and sRNAs were differentially expressed between strains D23580 and 4/74. A parallel proteomic approach confirmed that many of the gene expression differences led to alterations at the protein level. The differential expression of 199 genes and sRNAs within macrophages allowed us to predict functions of African *S.* Typhimurium ST313 that are modified during infection. The comparative gene expression data were used to predict key phenotypic differences between the pathovariants which are summarized in S1 Table. The power of our experimental approach is highlighted by our discovery of the molecular basis of the melibiose utilization defect of D23580, and a novel bacterial plasmid maintenance system that relied upon a plasmid-encoded essential gene.

In the future, functional transcriptomics could shed light on the factors responsible for the phenotypic differences that distinguish pathovariants of many bacterial pathogens.

## Materials and Methods

### Bacterial strains

The clinical isolate *Salmonella enterica* serovar Typhimurium D23580 was obtained from the Malawi-Liverpool-Wellcome Trust Clinical Research Programme, Blantyre, Malawi [19]. This strain, isolated from blood of an HIV- child from Malawi, is used as a representative of the *Salmonella* sequence-type ST313 after approval by the Malawian College of Medicine (COMREC ethics no. P.08/14/1614). *S.* Typhimurium 4/74 was originally isolated from the bowel of a calf with salmonellosis [61] and is used as a representative strain of *Salmonella* ST19. Other *Salmonella* strains referenced in this study are listed in S9 Table.

### Growth conditions

All strains were routinely grown in Lennox broth (LB) containing 10 g/L tryptone, 5 g/L yeast extract and 5 g/L NaCl. Liquid bacterial cultures were incubated at 37ºC 220 rpm for 16 h. Agar plates were prepared with 1.5% Bacto Agar (BD Difco). To test the ability to grow with melibiose as the sole carbon source, strains were grown in M9 minimal medium with 0.4% of melibiose. M9 minimal medium consisted of 1x M9 Minimal Salts (Sigma-Aldrich), 2 mM MgSO_4_, and 0.1 mM CaCl_2_. Glucose was added at a final concentration of 0.4% to M9 minimal medium to study growth behaviour of the *cysS* mutants. Media were supplemented with antibiotics when required: kanamycin (Km) 50 μg/mL, gentamicin (Gm) 20 μg/mL, tetracycline (Tc) 20 μg/mL, nalidixic acid (Nal) 50 μg/mL, and chloramphenicol (Cm) 20 μg/mL.

Details for growing bacteria in the sixteen *in vitro* infection-relevant conditions and inside murine RAW264.7 macrophages (ATCC TIB-71) have been published previously [37,39].

### Resequencing of *S.* Typhimurium D23580 genome

For PacBio sequencing, *S.* Typhimurium D23580 was grown for 16 h in Lennox medium at 37oC 220 rpm. DNA was extracted using the Bioline mini kit for DNA purification (Bioline). Genomic quality was assessed by electrophoresis in a 0.5% agarose gel at 30-35 V for 17-18 h. A 10 kb library was prepared for DNA sequencing using three SMRT cells on a PacBio RSI I (P5/C3 chemistry) at the Centre for Genomic Research, University of Liverpool, UK. Illumina sequencing of *S.* Typhimurium D23580 was performed by MicrobesNG, University of Birmingham, UK.

All the SNP and indel differences found between the chromosome and pSLT-BT sequences of the D23580 strain used in this study (accession: XXXXXXXXXX) and the published D23580 (accession: FN424405 and FN432031) were confirmed by PCR with external primers and subsequent Sanger sequencing.

Draft sequences of the pBT2 and pBT3 plasmids were provided by Robert A. Kingsley [19], and were used to design oligonucleotides for primer walking sequencing (all primer sequences are listed in S10 Table, Eurofins Genomics). Plasmid DNA from *S.* Typhimurium D23580 was isolated using the ISOLATE II Plasmid Mini Kit (Bioline). For pBT2, the following oligonucleotides were used: Fw-pBT2-1 and Rv-pBT2-1, Fw-pBT2-2 and Rv-pBT2-2; and, for pBT3, the following oligonucleotides were used: Fw-pBT3-3 and Rv-pBT3-3, Fw-pBT3-1 and Rv-pBT3-4, Fw-pBT3-4 and Rv-pBT3-2.

The resulting genome sequence was designed D23580_liv (accession: XXXXXXXXXX).

### Assembly of the *S.* Typhimurium D23580 complete genome

HGAP3 [62] was used for PacBio read assembly of the D23580 chromosome, and for the large plasmids pSLT-BT and pBT1. A hybrid assembly approach, Unicycler v0.4.5 [63], was used to combine the long reads from PacBio and the short reads from the Illumina platform in order to assemble small plasmids (not covered by PacBio due to size selection in library preparation) and to improve the large plasmids assemblies.

### RNA isolation, cDNA library preparation and Illumina sequencing

Total RNA from the sixteen *in vitro* growth conditions (EEP, MEP, LEP, ESP, LSP, 25ºC, NaCl, bile shock, low Fe^2+^ shock, anaerobic shock, anaerobic growth, oxygen shock, NonSPI2, InSPI2, peroxide shock, and nitric oxide shock) and murine RAW264.7 macrophages was isolated using TRIzol, and treated with DNase I, as described previously [37,39].

For RNA-seq, cDNA libraries were prepared and sequenced by Vertis Biotechnologie AG (Freising, Germany). Briefly, RNA samples were fragmented with ultrasound (4 pulses of 30 sec at 4°C), treated with antarctic phosphatase, and rephosphorylated with polynucleotide kinase (PNK). RNA fragments were poly(A)-tailed and an RNA adapter was ligated to the 5’-phosphate of the RNA. First-strand cDNA synthesis was carried out using an oligo(dT)-adapter primer and M-MLV reverse transcriptase. cDNA was subsequently amplified by PCR to 20-30 ng/μL, and purified using the Agencourt AMPure XP kit (Beckman Coulter Genomics). cDNA samples were pooled in equimolar amounts, size-selected to 150-500 bp and sequenced on an Illumina HiSeq 2000 system (single-end 100 bp reads). Minor changes were applied to different RNA-seq runs. For the third macrophage biological replicate of D23580, cDNA was PCR-amplified to 10-20 ng/μL, size-selected to 200-500 bp, and samples were sequenced on an Illumina HiSeq 2500 platform (1×100 bp). For RNA samples of D23580 and 4/74 grown in the four *in vitro* growth conditions with three biological replicates, the third macrophage replicate of 4/74, and the D23580 pBT1-cured strain, cDNA was PCR-amplified to 10-20 ng/μL and size-selected to 200-500 bp, and cDNA libraries were single-read sequenced on an Illumina NextSeq 500 system using 75 bp read length.

### Read processing and alignment

The quality of each RNA-seq library was assessed using FastQC v0.11.5 (http://www.bioinformatics.babraham.ac.uk/projects/fastqc/) and then processed with Trimmomatic v0.36 [64] to remove Illumina TruSeq adapter sequences, leading and trailing bases with a Phred quality score below 20 and trim reads with an average base quality score of 20 over a 4 bp sliding window. All reads less than 40 nucleotides in length after trimming were discarded from further analysis.

The remaining reads of each library were aligned to the corresponding genomes using Bowtie2 v2.2.9 [65] and alignments were filtered with Samtools v1.3.1 [66] using a MAPQ cut-off of 15. For *S.* Typhimurium D23580, reads were aligned to the sequences of the chromosome and the pSLT-BT, pBT1, pBT2, and pBT3 plasmids (accession: XXXXXXXXXX). For *S.* Typhimurium 4/74, reads were aligned to the sequences of the published 4/74 chromosome, and the plasmids pSLT^SL1344^, pCol1B9^SL1344^, and pRSF1010^SL1344^ (accession: CP002487, HE654724, HE654725, and HE654726, respectively). The RNA-seq mapping statistics are detailed in S4 Table. Reads were assigned to genomic features using featureCounts v1.5.1 [67].

The complete RNA-seq pipeline used for this study is described in https://github.com/will-rowe/rnaseq.

Two strain-specific browsers were generated for the visualization of the transcriptional data in a genomic context online (bioinf.gen.tcd.ie/cgi-bin/salcom_v2.pl). The different tracks in each JBrowse [50] were normalized using a published approach [68].

### Quantifying differences in expression with only one biological replicate

Expression levels were calculated as Transcripts per Million (TPM) values [40,41], and generated for coding genes and noncoding sRNAs in the chromosome, pSLT-BT and pBT1 plasmids for D23580 using the re-annotated D23580_liv genome (S2 Table). In 4/74, TPM values were determined for coding genes and noncoding sRNAs in the chromosome, and the three plasmids pSLT^4/74^, pCol1B9^4/74^, and pRSF1010^4/74^ [35]. Based on those values, and following previously described Materials and Methods, the expression cut-off was set as TPM > 10 for genes and sRNAs [37].

For comparative analysis between the two *S.* Typhimurium strains D23580 and 4/74, TPM values were obtained for the 4675 orthologous genes and noncoding sRNAs. These values were used to calculate fold-changes between strains. Due to the availability of only one biological replicate per growth condition, a conservative cut-off of ≥ 3 fold-change was used as a differential expression threshold between strains.

### Differential gene expression analysis with three biological replicates

Raw read counts from the 4674 orthologous coding genes and noncoding sRNAs for the three replicates of the five conditions (ESP, anaerobic growth, NonSPI2, InSPI2, macrophage) for 4/74 and D23580 were uploaded into Degust (S1 Data) (http://degust.erc.monash.edu/). Data were analysed using the Voom/Limma approach [69,70] with an FDR of ≤ 0.001 and Log_2_FC of ≥ 1. Pairwise comparisons were generated between the two strains for each specific condition. To remove genes with low counts across all samples, thresholds of ≥ 10 read counts and ≥ 1 Counts Per Million (CPM) in at least the three biological replicates of one sample were used [69,71].

For RNA-seq analysis of the D23580 ΔpBT1 strain grown in ESP, only two RNA-seq biological replicates were used for differential expression analysis with the three biological replicates of D23580 WT.

### Sample processing for proteomics

An LC-MS/MS (Q Exactive orbitrap) 4 h reversed phase C18 gradient was used to generate proteomic data from six biological replicates of each strain, 4/74 and D23580, grown in the ESP condition in Lennox broth. Pellet from bacterial cultures was resuspended in 50 mM phosphate buffer (pH 8), sonicated (10 sec ON, 50 sec OFF, for 10 cycles at 30% amplitude), and supernatants were analysed after centrifugation at 16,000 g for 20 min. Subsequent experimental procedures were performed at the Centre for Proteome Research at the University of Liverpool, UK. In brief, 100 μg of protein were digested (RapiGest, in-solution trypsin digestion) and 1 μg of digested protein was run on an LC-MS/MS platform.

### Analysis of proteomic data

A database was generated merging the amino acid sequences of the annotated genes in 4/74 [35] and our re-annotated D23580 to allow homologous proteins as well as strain-specific proteins to be identified. The merged database was clustered using the program Cd-hit and an identity threshold of 95% [72]. Clusters with a single protein, representing strain-specific proteins, were included in the database with their accession ID. Clusters with more than one protein represented orthologues, and only peptides common to all proteins of the cluster were included in the database. Common peptides allowed label-free comparison of proteins that had a low level of sequence variation.

Raw data obtained from the LC-MS/MS platform were loaded into the Progenesis QI software (Nonlinear Dynamics) for label-free quantification analysis. Differential expression analysis between the two strains, 4/74 and D23580, is shown in S7 Table. From those results (2013 proteins), multihit proteins (peptides assigned to more than one protein in the same strain) were removed leaving a total of 2004 proteins. Cut-offs of ≥ 2 unique peptides *per* identified protein (1632 proteins), ≥ 2 fold-change expression and *p*-value < 0.05 between strains (121 proteins) were used. Among the 121 proteins, 25 were 4/74-specific, 30 were D23580-specific, and 66 were encoded by orthologous genes between strains.

### Alpha-galactosidase activity

To assess alpha-galactosidase activity, strains were grown on Pinnacle™ *Salmonella* ABC (chromogenic *Salmonella* medium, LabM). Bacteria that are able to produce alpha-galactosidase in the absence of beta-galactosidase appear as green colonies on this medium due to the hydrolysis of X-alpha-Gal. This enzymatic activity was correlated to the ability to grow in M9 minimal medium with melibiose as the sole carbon source.

### Construction of scarless single-nucleotide substitution mutants

Two strategies were used for single-nucleotide replacement as previously described [20]. For D23580 *flhA^4/74^*, a single-strand DNA oligonucleotide recombination approach was used [73]. Briefly, the flhA-474SNP oligonucleotide containing the SNP in 4/74 (‘C’) was used to replace the SNP in D23580 (T). The methodology followed the same strategy used for λ Red recombination explained below. After electroporation of the ssDNA oligonucleotide into D23580 carrying the pSIM5-*tet* plasmid, screening for D23580 recombinants was performed using a PCR with a stringent annealing temperature and primers Fw-flhA and Rv-flhA-474SNP. The reverse primer contained the 4/74 SNP in *flhA*. The SNP mutation in D23580 *flhA^4/74^* was confirmed by Illumina whole-genome sequence (MicrobesNG, University of Birmingham). Variant-calling bioinformatic analysis confirmed the intended mutation and the absence of secondary nonintended mutations.

The second strategy for constructing scarless SNP mutants followed a previously described approach based on the pEMG suicide plasmid [24,74]. Oligonucleotides melR-EcoRI-F and melR-BamHI-R were used to PCR-amplify, in 4/74 and D23580, a *melR* region containing the SNP described between strains. Additionally, primers melB-EcoRI-F and melB-BamHI-R were used for amplification, in 4/74 and D23580, of a *melB* region containing the two SNPs described in this gene. PCR products were cloned into the pEMG suicide plasmid and transformed into *E. coli* S17-1 *λpir*. The resulting recombinant plasmids were conjugated into 4/74 or D23580, depending on the strain that was used for the PCR-amplification. For *S.* Typhimurium 4/74, transconjugants were selected on M9 minimal medium with 0.2% glucose and Km. For *S.* Typhimurium D23580, transconjugants were selected on LB Cm Km plates. As described previously [24], transconjugants were transformed with the pSW-2 plasmid to promote the loss of the integrated pEMG by a second homologous recombination. The single-nucleotide substitutions were confirmed by PCR amplification with external primers and sequencing. Mutants D23580 *melR^+^* and D23580 *melR^+^melB^+^* were confirmed by Illumina whole-genome sequencing (MicrobesNG, University of Birmingham). Variant-calling bioinformatic analysis confirmed the intended mutations and the absence of secondary nonintended mutations in D23580 *melR^+^melB^+^*. Mutant D23580 *melR^+^* had a secondary nonintended synonymous mutation at the chromosomal location 436,081 in *STMMW_04211* (GCC → GCA).

### Construction of the Δ*cysSc^hr^* and Δ*cysS^pBT1^* mutants in *S.* Typhimurium D23580 by λ Red recombineering

The D23580 mutants in *cysS^chr^ (STMMW_06051*) and *cysS^pBT1^ (pBT1-0241*) were constructed using the λ Red recombination strategy [75]. The kanamycin resistance cassette (*aph*) of pKD4 was amplified by PCR using the primer pairs NW_206/NW_207 and NW_210/NW_211, respectively. The resulting PCR fragments were electroporated into D23580 carrying the recombineering plasmid pSIM5-*tet* following the previously described methodology [20,76]. The *ΔcysS^chr^::aph* mutation was transduced into D23580 wild type using the high frequency transducing bacteriophage P22 HT 105/1 *int-201* [77] as previously described [24]. The D23580 *ΔcysSp^BT1^*::*aph* mutant was whole-genome sequenced using the Illumina technology (MicrobesNG, University of Birmingham). Variant-calling bioinformatic analysis confirmed the intended mutation and the absence of secondary nonintended mutations with the exception of a six-nucleotide insertion in a noncoding region at the chromosomal position 2,755,248 (A → AGCAAGG). The Km resistance cassettes of the two recombinant strains, *ΔcysS^chr^::aph* and D23580 *ΔcysSp^BT1^::aph*, were flipped-out using the FLP recombinase expression plasmid pCP20-TcR [22].

### Construction of the *S.* Typhimurium D23580 pBT1-cured strain

The pBT1 plasmid was cured from D23580 using published methodology [78]. First, the *pBT1-0211* gene of pBT1, encoding a putative RelE/StbE replicon stabilization toxin, was replaced by a *I-SceI-aph* module by λ Red recombination. The *I-SceI-aph* module was amplified from pKD4-I-SceI [24] using primers NW_163 and NW_164, and the resulting PCR fragment was electroporated into D23580 carrying pSIM5-*tet*. The resulting *ΔpBT1-0211::I-SceI-aph* mutants were selected on LB Km plates and the mutation was transduced into D23580 wild type as described above. D23580 *ΔpBT1-0211::I-SceI-aph* was subsequently transformed with the I-SceI meganuclease producing plasmid pSW-2 [74], and transformants were selected on LB Gm agar plates supplemented with 1 mM *m*-toluate, which induces high expression of the I-SceI nuclease from pSW-2. The absence of pBT1 was confirmed by whole-genome sequencing of the D23580 ΔpBT1 strain (MicrobesNG, University of Birmingham).

### Growth curves in M9 melibiose, LB and M9 glucose media

Overnight bacterial cultures were washed twice with PBS and resuspended in the specific growth medium at an O.D._600 nm_ of 0.01. Growth curves of strains grown in M9 minimal medium supplemented with melibiose or glucose were based on O.D. at 600 nm measurements every hour of samples growing in a 96-well plate. Microplates were incubated at 37ºC on an orbital shaker set at 500 rpm in a FLUOstar Omega (BMG Labtech) plate reader. Only the values of the O.D._600 nm_ at 8 h were plotted for strains grown in M9 melibiose medium. For the O.D._600 nm_ measurement of D23580, D23580Δ*cysS^chr^*::*frt*, D23580 *ΔcysSp^BT1^::frt* and D23580 ΔpBT1 grown in LB, an Infinite F Nano+ (Tecan) plate reader was used with an orbital shaking of 432 rpm.

### Analysis of SNP conservation in the melibiose utilization operon

The conservation of the two SNPs in *melB* and one SNP in *melR*, that distinguished the *S.* Typhimurium strains D23580 and 4/74, was analysed in the genomes of 258 *S.* Typhimurium ST313 isolates from Malawi and the United Kingdom. The A5 assembly pipeline [79] and ABACAS [80] were used when a reference quality genome was not available. The PanSeq package allowed the identification of core genome SNPs [81] and the concatenated SNP alignment served to obtain a maximum-likelihood phylogenetic tree using PhylML [82]. BLASTn was used to identify the genotype of the melibiose SNPs shown in Fig 6C in all genomes (S8 Table).

### Conservation of pBT1 and pSLT-BT plasmids among ST313 isolates

For phylogenetic analysis of ST313 isolates, all available FASTQ data was downloaded from the ENA using FASTQ dump v2.8.2 (accessions in S8 Table, ENA access date: 01.02.2017). Data quality was assessed using FastQC v0.11.5 (http://www.bioinformatics.babraham.ac.uk/projects/fastqc/) and then processed with Trimmomatic v0.36 [64] to any adapter sequences, leading and trailing bases with a Phred quality score below 20 and trim reads with an average base quality score of 20 over a 4 bp sliding window. All reads less than 40 nucleotides in length after trimming were discarded from further analysis.

A multiple sequence alignment was generated by mapping isolate FASTQ data to the ST313 D23580 reference genome (pSLT-BT and pBT1 plasmids) (accession: XXXXXXXXXX) using Bowtie2 v2.2.9 [65]. Alignments were filtered (MAPQ cut-off 15) and then deduplicated, sorted and variant called with Samtools v1.3.1 [66]. For each alignment, recombination was masked using Gubbins v2.2.0 [83] and the variable sites were used to construct a maximum likelihood tree using RAxML [84]. Phylogenetic trees were visualised using Figtree (http://tree.bio.ed.ac.uk/software/figtree/) and Dendroscope [85]. Coverage information was extracted from the alignment files using bedtools v2.26.0 [86] and visualised using R. Results are shown in Fig 8D (S8 Table).

### Statistical analysis for phenotypic studies

Unpaired two-tailed Student’s t-tests were performed using GraphPad Prism 6.0 (GraphPad Software Inc., La Jolla, CA, USA).

### Data availability

The updated D23580 genome and annotation (D23580_liv) have been deposited in the European Nucleotide Archive (ENA) repository (EMBL-EBI) under accession XXXXXXXXXX. The strain has been deposited in the National Collection of Type Cultures (NCTC) from Public Health England under accession number: pending.

The RNA-seq-derived transcriptomic data generated and re-analysed in this study have been deposited in the Gene Expression Omnibus (GEO) database: accession number YYYYYYYYY.

The mass spectrometry proteomics data have been deposited to the ProteomeXchange Consortium via the PRIDE [87] partner repository with the dataset identifier ZZZZZZZZZ.

Resources for the visualization of the RNA-seq data in the 16 *in vitro* growth conditions and the intra-macrophage environment are available online at bioinf.gen.tcd.ie/cgi-bin/salcom_v2.pl.

## Acknowledgments

We are grateful to present and former members of the Hinton laboratory for helpful discussions, particularly Aoife Colgan and Sathesh Sivasankaran, and Paul Loughnane for his expert technical assistance. We appreciated Nicholas Feasey’s insightful comments during this project. We thank Margaret Hughes, and John Kenny from the Centre for Genomic Research at the University of Liverpool for technical assistance for PacBio sequencing, and Alistair Darby for many helpful discussions. We appreciate the assistance and discussions provided by Lynn McLean and Robert Beynon from the Centre for Proteome Research at the University of Liverpool. We are grateful to Fritz Thϋmmler (Vertis Biotechnologie AG) for the construction of high-quality cDNA libraries. We also thank Gemma Langridge and Nick Thomson for their help with improving the D23580 annotation.

## Author contributions

R.C. and J.C.D.H. designed the study. R.C., D.L.H., C.K., W.Y.F., L.L.-L., X.Z., N.W., and S.E.C. performed wet-lab experiments. R.C., S.V.O., P.B., R.R.C., W.P.M.R., A.V.P. analysed data. R.C. and J.C.D.H. interpreted the data. J.H., D.M.M., and M.A.G. contributed to data interpretation. R.A.K. provided draft sequences for pBT1, pBT2, and pBT3. K.H. created the *SalComD23580* website. R.C. and J.C.D.H. wrote the manuscript.

## Supporting Information

**S1 Table**. Phenotypic features that distinguish *S.* Typhimurium ST313 from ST19 isolates from the literature.

**S1 Text**. Supporting Materials and Methods.

**S1 Fig**. Schematic representation of the RNA-seq-based comparative transcriptomic approach.

**S2 Table**. Complete *S.* Typhimurium D23580 updated annotation with 4/74 orthologies.

**S3 Table**. SNPs, MNPs, and indels between *S.* Typhimurium 4/74 and D23580.

**S2 Fig. Novel *S.* Typhimurium D23580 noncoding sRNAs**. Northern blots confirming the existence of novel sRNAs annotated in the BTP1 prophage region (S1 Text). For every individual sRNA, a northern blot and mapped reads in the same conditions are shown. The arrowheads indicate the most prominent bands. 5S rRNA was used as a loading control. Estimated length of the sRNAs is in brackets and was based on RNA-seq data and sequence analysis. Transcription start sites (TSS) are highlighted at the bottom.

**S4 Table. RNA-seq sequence reads for *S.* Typhimurium 4/74 and D23580**.

**S5 Table. TPM values for *S.* Typhimurium 4/74 and D23580 from the two RNA-seq datasets**.

**S3 Fig. Transcriptional response to infection-relevant stress of *S.* Typhimurium 4/74 and D23580.** (A) Percentage of expressed genes (TPM > 10) for each individual strain. (B) Number of coding genes and sRNAs differentially expressed (fold-change ≥ 3) for each of the 17 infection-relevant conditions. (C) Heat map of the cluster analysis of all orthologous coding genes and sRNAs between the two strains obtained using GeneSpring GX7.3 (Agilent). The transcriptional expression value (TPM) for each coding gene and sRNA in each condition in D23580 was divided by the TPM value for the same gene/sRNA and condition in 4/74. (D) Bubble chart for 4/74 representing up-regulated coding genes and sRNAs versus down-regulated. The following comparisons based on TPM values were obtained for each specific condition: MEP, LEP, ESP, and LSP were compared to EEP; NaCl shock, bile shock, low Fe^2+^ shock, and anaerobic shock were compared to MEP; oxygen shock was compared to anaerobic growth; peroxide shock and nitric oxide shocks were compared to InSPI2; InSPI2 was compared to NonSPI2; and macrophage was compared to ESP. (E) Bubble chart for D23580.

**S4 Fig. The Tn*21*-like antibiotic resistance cassette is inserted in the *mig-5* operon preventing expression of *rlgAb, rlgAa*, and *pSLT043***. Visualization of the RNA-seq data in the 17 infection-relevant conditions from the online JBrowse resources provided in this study. Red arrows represent genes that showed upregulation in 4/74 versus D23580, and blue arrows represented D23580-downregulated genes. Scale of the mapped reads was 1 to 500. The insertion of the Tn*21*-like element is indicated by dotted lines.

**S5 Fig. Differences in expression of the flagellar regulon between *S.* Typhimurium 4/74 and D23580**. (A) Heat map of the flagellar regulon genes representing the relative expression of D23580 versus 4/74 in five growth conditions. TPM values were obtained from the RNA-seq dataset with only one biological replicate. (B) Swimming motility assay of 4/74, D23580, and D23580 *flhA^4/74^* (S1 Text). Bars represent the mean of 12 independent replicates and standard deviation. Significant differences indicate ****, *p*-value < 0.0001; and **, *p*-value < 0.01. (C) Lactate dehydrogenase (LDH) cytotoxicity assay using BMDM C57BL/6 macrophages (S1 Text). Bars represent the mean of 6 independent replicates and standard deviation. Groups were compared using one-way Anova and Tukey’s multiple comparisons test, significant differences indicate ****, *p*-value < 0.0001; and **, *p*-value < 0.01. (D) Heat map of the flagellar regulon using the RNA-seq data with three biological replicates. Results represent the fold-change (D23580 versus 4/74) and FDR values obtained from Degust.

**S6 Table. RNA-seq results for *S.* Typhimurium 4/74 and D23580 from Degust**.

**S6 Fig. Virulence-associated genes differentially-expressed between *S.* Typhimurium 4/74 and D23580**. (A) Heat map of the *Salmonella* pathogenicity islands genes and sRNAs that show ≥ 2 fold-change and ≤ 0.001 FDR (D23580 versus 4/74), obtained using GeneSpring GX7.3 (Agilent). The CPM values of three biological replicates for each coding gene and sRNA in each condition in D23580 were compared to the CPM values for the same gene/sRNA and condition in 4/74. (B) Differential gene expression of D23580 versus 4/74 in the early stationary phase (on the left) and the macrophage (on the right) conditions. Colors refer to fold-changes of D23580 versus 4/74 from differential expression analysis using Degust, red = D23580-upregulated, blue = D23580-downregulated. The figure includes genes that are differentially expressed ≥ 4 fold-change with ≤ 0.001 FDR (red and blue font color). Purple and light blue font colors represent up-regulated or down-regulated genes, respectively, that are related to the previous functional groups of genes, but have a fold-change ≤ 4 and ≥ 2 (≤ 0.001 FDR).

**S7 Fig. Reproducibility of transcriptomic experiments**. Correlation coefficient plots of Log_2_[TPM values] for 5 infection-relevant conditions. The “different sequencing runs” plots compare the two RNA-seq datasets (one biological replicate versus three biological replicates). The “same sequencing run” plots compare two samples of the RNA-seq dataset with three biological replicates. (A) 4/74, (B) D23580.

**S7 Table. Proteomic data of *S.* Typhimurium 4/74 and D23580 grown in ESP**.

**S8 Table. Conservation of the SNPs in the melibiose operon and the pBT1 and pSLT-BT plasmids among *S.* Typhimurium ST313 isolates**.

**S9 Table. Bacterial strains and plasmids**.

**S10 Table. Oligonucleotides used in this study**.

**S1 Data. Input file for Degust**. Data include counts for the three biological replicates in five growth conditions (ESP, anaerobic growth, NonSPI2, InSPI2, macrophage) in 4/74 and D23580, and for the two biological replicates of D23580 ΔpBT1 in ESP.

